# New algorithms for unsupervised cell clustering from scRNA-seq data

**DOI:** 10.1101/2024.11.22.624768

**Authors:** Melissa Robles, Jorge Díaz-Riaño, Cristhian Forigua, Sebastian Ojeda, Laura Guio, Paula Siaucho, Jennifer Guzman-Porras, Danilo García-Orjuela, Andres Naranjo, Silvia Maradei, Adolfo Quiroz, Jorge Duitama

## Abstract

The identification of cell types is a basic step of the pipeline for Single-Cell RNA sequencing data analysis. However, unsupervised clustering of cells from scRNA-seq data has multiple challenges: the high dimensional nature of the data, the sparse nature of the gene expression matrix, and the presence of technical noise that can introduce false zero entries. In this study, we introduce new algorithms for clustering scRNA-seq data. The first algorithm builds a ***k***-MST graph from distances obtained directly from the input data without dimensionality reduction. The computation follows an iterative procedure of k steps in which each step calculates and stores the edges of minimum spanning trees over different subgraphs obtained removing edges selected in previous iterations. The Louvain algorithm is executed on the ***k***-MST graph for cell clustering. We also explored alternatives based on neural networks in which an autoencoder is used to learn the parameters of a Gaussian mixture model, aiming to improve the handling of clusters with different shapes and sizes. Benchmark experiments with simulated data and public datasets show that the algorithms proposed in this work have competitive accuracy, compared to previous solutions, but also that sequencing depth, number of cells and tissue types have important effects on the performance of the algorithms. Moreover, we performed further experiments with scRNA-data taken from a patient with refractory epilepsy. The AE-GMM model achieved the best accuracy for this dataset, and the ***k***-MST ranked first among methods that do not require previous information on the expected number of clusters.

## 1 Introduction

The Single Cell RNA sequencing (scRNA-seq) methodology enables the quantification of gene expression at a single cell level. Since the initial proof of concept in 2009 [1] the use of this technology has increased steadily, thanks to different improvements in cell sorting and library preparation techniques, and the reduction of sequencing costs [2]. Gene expression profiles based on scRNA-seq data can be used to differentiate according to cell type, which refers to the category or classification of cells based on their specific characteristics, such as molecular composition, morphology, or function [3]. Aggregating the data by cell types facilitates the identification of genes whose expression significantly differs among the various cell types present in a tissue. Additionally, it improves the identification of genes whose expression varies between tissues affected by diseases and healthy tissues, compared to bulk RNA-seq data.

Identifying cell types is a primary step of the data analysis pipeline for Single-Cell sequencing experiments. However, unsupervised clustering of scRNA-seq data has multiple challenges. One challenge is the high-dimensional nature of the data, where the number of features (genes in this case) can be in the thousands. A large number of features tends to homogenize the distances between samples, which can reduce the accuracy of different clustering algorithms [4, 5]. Another challenge is the sparse nature of the gene expression matrix. Not all genes are active in every cell, leading to many zero entries, often representing more than 90% of the data to analyze. The final challenge is the presence of technical noise that can introduce false zero entries, known as “dropouts”.

Multiple methodologies have been proposed for cell type detection in scRNA-seq data. A combination of classical techniques for unsupervised clustering is implemented in the SC3 library [6]. SC3 starts with the computation of three distance matrices: Euclidean distance, Spearman correlation, and Pearson correlation, followed by dimensionality reduction of these matrices through principal component analysis (PCA). Subsequently, *k*-means clustering is applied to the three resulting matrices, and finally, hierarchical clustering is performed using the consensus of the three obtained results. Some methodologies, such as SINCERA [7] and DP-SIMLR [8], are based on hierarchical and density clustering algorithms.

Graph-based approaches are also used as cell clustering methodologies for scRNA data. An example of this approach is the SSNN-Louvain methodology [9], which relies on community detection with a modified Louvain algorithm on the Shared-Nearest-Neighbor (SNN) graph. A graph-based approach is also implemented in the widely used tool Seurat [10]. This software employs community detection from the graph of *k* nearest neighbors (*k*-NN) of the data obtained after dimensionality reduction using PCA. Although various methods can be applied to obtain the communities, the default approach of Seurat is the Louvain algorithm [11]. This algorithm optimizes a modularity function, looking for dense connections between nodes within communities, and sparse connections between nodes of different communities. The SLM algorithm [12] is an alternative technique to optimize the modularity, available in Seurat. In contrast to the Louvain algorithm, SLM allows the movement of entire sets of nodes and the splitting of communities, providing greater flexibility in exploring solutions for modularity optimization. Constructing the graph itself is particularly crucial when using community detection methods in graphs. Seurat constructs the initial graph from the *k* nearest neighbors graph based on Euclidean distance and defines the weights of the resulting edges as the Jaccard similarity.

Neural networks have also been proposed as alternatives for analysis of scRNA data. In particular, an autoencoder is a type of neural network that can serve to a dual purpose of error correction and dimensionality reduction [13, 14]. The models learn to encode data into a lower-dimensional space and then reconstruct it by calculating a loss function that penalizes the differences between the original and reconstructed data. For scRNA-seq data, the use of autoencoders have shown positive results in both data clustering [15, 16] and data imputation [17, 18]. Following the approach proposed by Tian et al. [16], the encoder architecture can be modified to enable clustering during training, allowing it to learn the parameters of clusters distributions.

In this article, we introduce two novel clustering algorithms for scRNA-seq data. The first algorithm constructs a *k* minimum spanning tree (*k*-MST) graph using pairwise Pearson correlation between cells. Unlike other methods [6, 10], it omits PCA dimensionality reduction and instead selects the genes with the highest upward deviation from variance, adjusted by the mean of the counts. Following this, the Louvain algorithm is applied to identify clusters in the data. The second approach utilizes deep learning, where cluster parameters are learned in a secondary training stage with a GMM-log likelihood loss function. We assessed the accuracy of these methods against previous approaches using both simulated and real datasets, including a novel dataset taken from a patient with refractory epilepsy.

## 2 Algorithms

### 2.1 Clustering based on the *k*-MST graph

Figure 1 shows the main steps of the proposed *k*-MST algorithm for analysis of scRNA data. Reads obtained by the experiment (Figure 1a) are processed using one of the standard pipelines to generate the input count matrix, which contains the expression levels of each gene across all cells in the experiment (Figure 1b). During the preprocessing step, genes and samples are filtered using statistics on the expression levels per gene and per cell (Figure 1c, See methods 3.3 for details). Starting from the filtered matrix, the algorithm computes the Pearson’s correlation function over each pair of cells, building a similarity score matrix of dimensions *n × n* with *n* representing the number of cells in the dataset (Figure 1d): for each pair of gene expression vectors *x*_*i*_ and *x*_*j*_ corresponding to two different cells, the correlation is defined as:

**Fig. 1:**
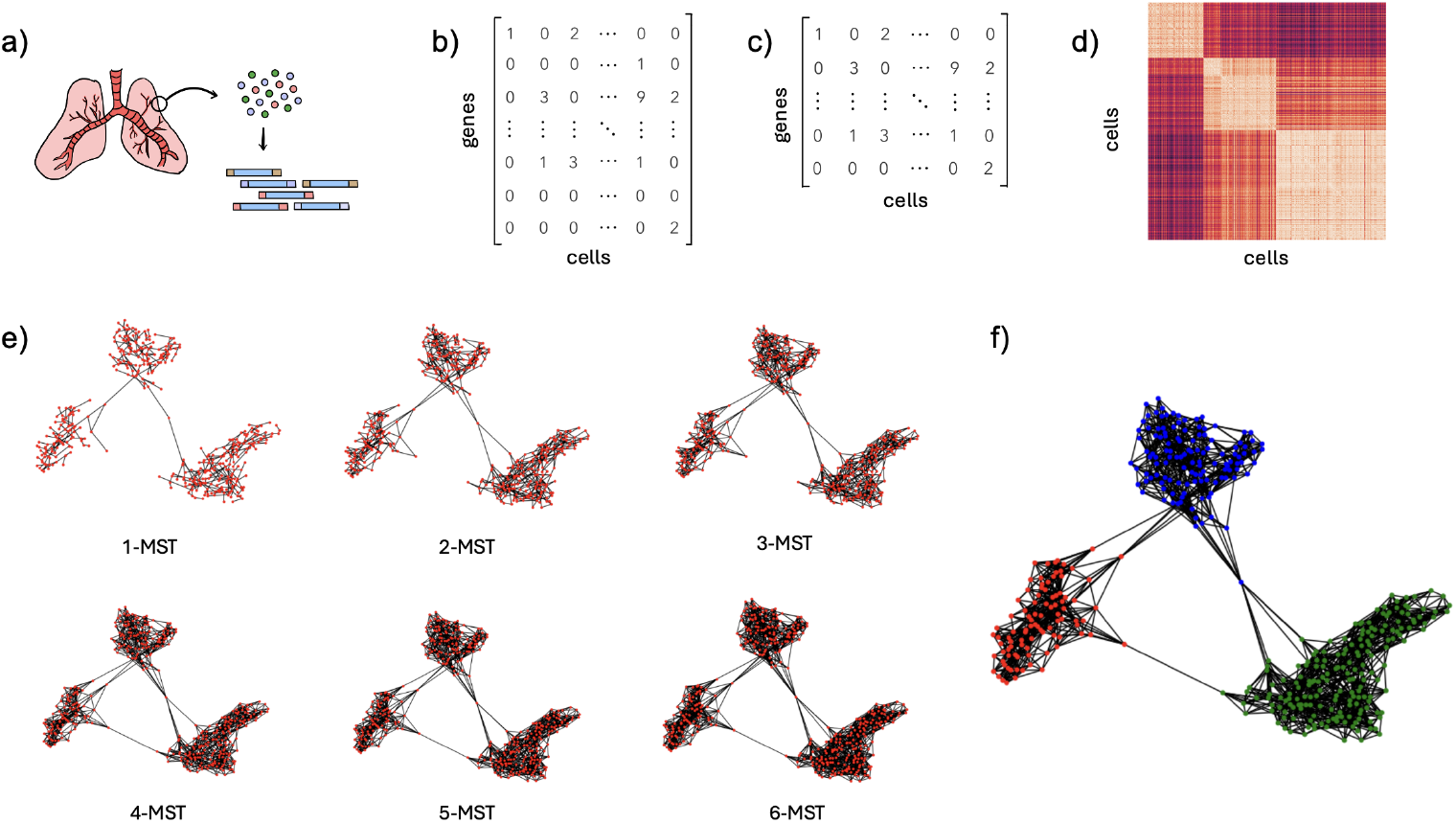
Graph-based scRNA-seq clustering pipeline. a) Perform single-cell RNA sequencing. b) Generate the count matrix. c) Gene filtering: Retain only the 5000 genes with the highest upward deviation from the adjusted variance. d) Compute the pairwise Pearson correlation between cells. e) Compute the k-MST graph f) Apply the Louvain algorithm for community detection.

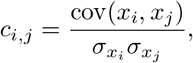

where cov(*x*_*i*_, *x*_*j*_) denotes the covariance between *x*_*i*_ and *x*_*j*_, and 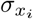 and 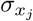 are the standard deviations of *x*_*i*_ and *x*_*j*_, respectively. The similarity matrix is the input of the main step of the algorithm, which is the construction of the *k*-MST graph (Figure 1e). This graph was first introduced in the context of the multivariate two-sample problem by Friedman and Rafsky in 1979 [19]. The computation follows an iterative procedure. Initially, the MST of the original graph is calculated and their edges are selected and removed from the original graph. A second MST is calculated on the resulting graph and the edges are selected and removed again. This iterative process continues for *k* = log(*n*) iterations, and the final graph is the union of the *k* “orthogonal” MST graphs.

Using the resulting graph, we use the Louvain algorithm [11] to partition the nodes, with the goal of optimizing the modularity of a graph *G* and a partition *C* defined as

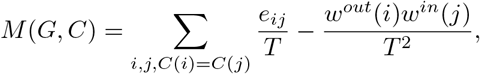

where the value *e*_*ij*_ corresponds to the weight of the edge (*i, j*), *T* represents the sum of all weight in the graph, *w*^*out*^(*i*) = ∑_*k*_ *e*_*ik*_ and *w*^*in*^(*j*) = _*k*_ *e*_*kj*_.

The Louvain algorithm begins by assigning each node to its own community. Through a series of iterations, nodes are reassigned to different communities to maximize modularity. This process continues until no further improvement in modularity can be achieved, resulting in the identification of stable community structures within the network (Figure 1f).

### 2.2 Clustering based on deep learning

Taking into account recent works in which autoencoder-based deep learning methods were used to perform dimensionality reduction and further clustering, we also explored alternatives of analysis following this approach. Our method combines the zero-inflated negative binomial (ZINB) loss with a loss function based on a Generalized Mixed Model (GMM) to improve the computation of clusters. Unlike previous approaches, which use *k*-Means based clustering assuming spherical clusters, the GMM-based clustering provides higher flexibility to capture clusters of various shapes, especially those that are ellipsoidal. The GMM-based loss function is based on the probability distribution of cells over the clusters. After dimensionality reduction, the model assumes that each cluster follows a Gaussian distribution with mean vector *μ* and covariance matrix Σ. The data likelihood function is modeled as follows:

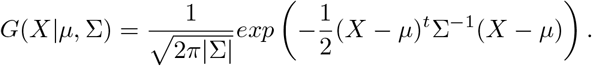

The training of this network is performed in two stages. First, the autoencoder is pretrained with the *ℒ*_*ZINB*_ loss function. Then, a fine-tuning stage is performed in which the latent representation is optimized, guided by the GMM loss function. The model learns the parameters of each Gaussian function at this stage to perform probabilistic clustering. Specifically, it learns three parameters: *π*_*c*_ *∈* ℝ, *μ*_*c*_ *∈* ℝ^32^, and Σ_*c*_ *∈* ℝ^32×32^ for each cluster *c*. The value of *μ*_*c*_ represents the mean of the cluster *c*. These mean values are initialized as the centroids found by the *k*-Means algorithm. For each cluster *c*, Σ_*c*_ represents the covariance matrix of *c* and *π*_*c*_ represents its weight, which can be understood as the prior probability that a cell is assigned to such cluster. Hence, *π*_*c*_ values are constrained to add upto 1:

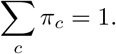

The *ℒ*_GMM_ loss is defined as:

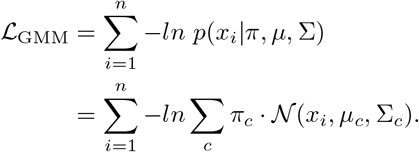

By incorporating this loss function, the model learns not only the mean, dispersion, and dropout as in DCA [15] but also the parameters that define the distributions of each data cluster, as illustrated in Figure 2. These data can be used to assign a probability to each point belonging to each cluster, also known as *soft clustering*.

**Fig. 2:**
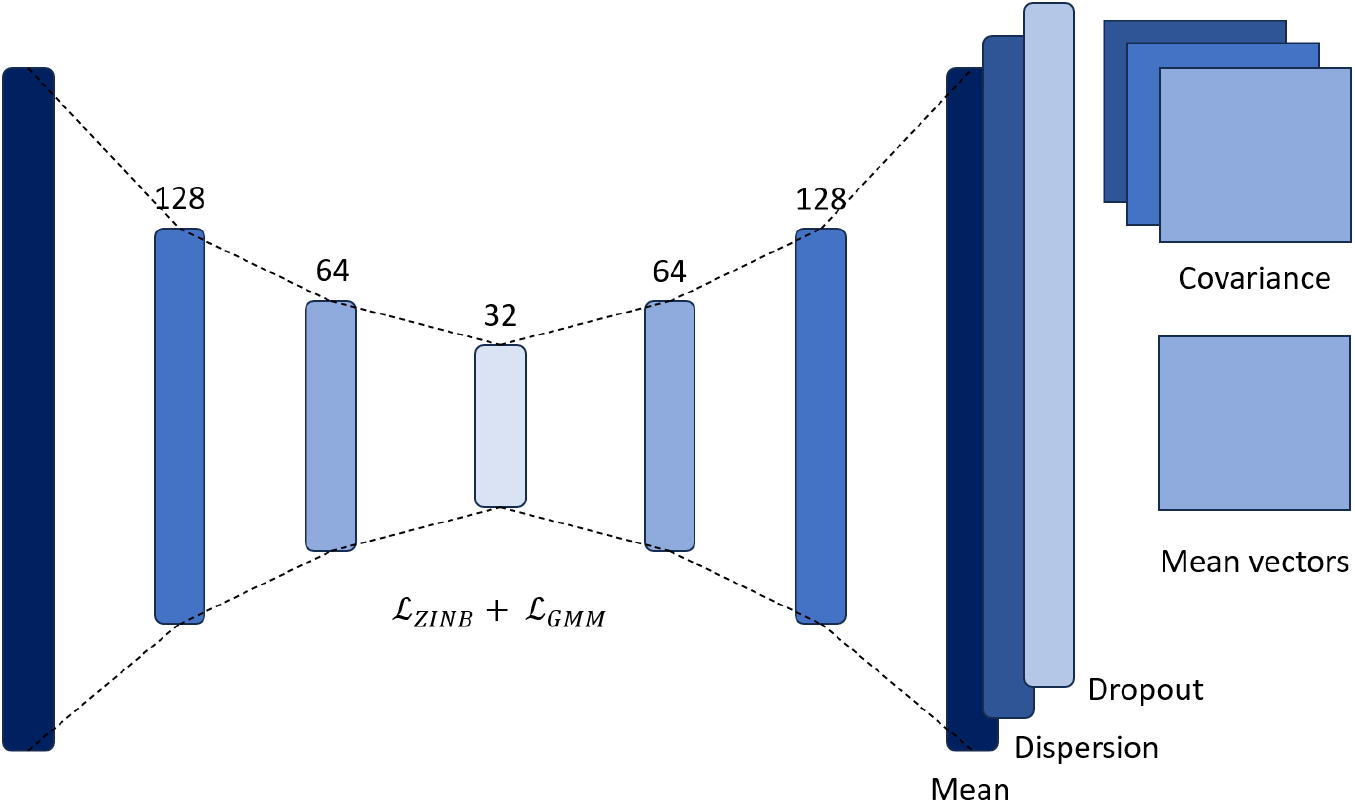
Neural network architecture. In the fine-tuning stage the model learns the parameters of each cluster.

## 3 Methods

### 3.1 Simulated data

We conducted simulations of scRNA-seq data to assess the performance of the proposed methodologies, using the SymSim tool [20]. This tool is specifically designed to model the processes underlying scRNA-seq experiments. It generates data by incorporating three primary sources of variation: intrinsic noise, extrinsic variation representing different cell states, and technical variation due to low sensitivity, measurement noise, and bias. This testing process included 24 different experiments varying the number of cells (500, 750, and 1000) the number of genes (1500 and 2000), the number of clusters (3 and 5), and the mean depth (1000 and 2000). In all experiments the minimum population size was set to 40 and the mean alpha was set to 0.01. We created five replicates for each experiment to assess the stability of the algorithms. The designed experiments had an average of 83.39% of zero entries with a standard deviation of 3.21% across replicates.

### 3.2 Non simulated data

We used five publicly available scRNA-seq datasets obtained from various species, tissues, and sample sizes for further accuracy assessment of the proposed methods (Table 1). Additionally, we sequenced a new sample taken from the brain of a patient with refractory epilepsy during a scheduled surgery. For public datasets, the clusters identified in the original studies served as gold standard labels.

**Table 1:** Comparison of the datasets by species, tissue, number of cells and the percentage of zeros in the data.

| Dataset | Species | Tissue | Cells | % Zeros |
| --- | --- | --- | --- | --- |
| 10X PBMC [21] | Human | Blood | 4271 | 96.08% |
| Human Liver [22] | Human | Liver | 8444 | 90.76% |
| Mouse bladder [23] | Mouse | Bladder | 2746 | 94.86% |
| Macosko mouse retina [24] | Mouse | Retina | 14653 | 88.34% |
| Worm neuron [25] | Worm | Neuron | 4186 | 98.61% |
| Epilepsy | Human | Brain | 4310 | 91.29% |

The epilepsy sample was selected from a cohort of patients treated by the Epilepsy Surgery service at Fundación Hospital de la Misericordia (HOMI) in Bogotá, Colombia. Patients diagnosed with refractory epilepsy, according to ILAE criteria, and identified as potential candidates for surgery by the pediatric epilepsy surgery group at HOMI were considered for inclusion. Exclusion criteria included autoimmune diseases, diabetes mellitus, and cancer. Participants who agreed to participate in the research study provided informed consent or assent. One patient with a diagnosis of right frontal structural and drug-resistant focal epilepsy was selected.

Brain tissue sample was collected from surgery, immediately transported to the laboratory on dry ice within 30 minutes of collection, and stored at −80°C until further processing. The frozen tissue was received and processed at the Princess Margaret Genomic Centre in Canada, according to the established protocols for nuclei extraction and purification of the sequencing center. The study employed the 10X Genomics 3’ v3.1 Nuc-Seq methodology, capturing roughly 7500 nuclei into droplets via the 10X Chromium system (3’ v3.1 kit). The library was designed to target an average read depth of 150,000 reads per nucleus. Mapping and feature counting were performed using STARsolo v2.7.11a [26] against the GRCh38-2020-A (Gencode v32 / Ensembl 98) human reference genome. The count matrix was then imported into Seurat v5 [27] for preprocessing. Cells with abnormal mitochondrial content (major than 5%) and cells with fewer than 100 or more than 10,000 expressed genes were removed. Doublets were identified and removed from the dataset using scDblFinder v3.15 package [28].

To use this dataset for benchmarking purposes, a gold standard set of labels was constructed using the single cell resource published under the Human Protein Atlas Project [29, 30]. This dataset comprises gene expression data across various human tissues and cell types, utilizing single-cell RNA sequencing (scRNA-seq), cell sorting, single-nuclei RNA sequencing (snRNA-seq), and bulk RNA-seq correlation analyses. It includes 557 individual cell type clusters representing 81 distinct cell types. Each cell was classified as neuronal (Inhibitory/GABAergic or Excitatory/Glutamatergic), non-neuronal (Astrocytes, Microglial cells, Oligodendrocyte precursor cells, Oligodendrocytes), or non-brain related cellular type (Others). For each cell in the dataset, a correlation matrix was computed between the gene expression profiles (nTPM) of the HPA reference. The cell type associated with the maximum correlation value for each cell was initially assigned as the biological classification. Finally, cells clustering near a group with a biological function different from their assigned annotation were manually removed from the dataset.

### 3.3 Preprocessing of the count matrices

We performed normalization in these experiments using the Scanpy package [31], which normalizes each cell by total counts over all genes, so that every cell has the same total count after normalization. Size factors are computed for each cell, by dividing the count sum of the specific cell across all genes, divided by a constant factor *L*. In the predefined parameters of Scanpy, *L* is equal to the median raw count depth in the dataset. The normalization is then followed by a logarithmic scaling to complete the processing of the count matrix. In the Neural Network, an additional step was carried out following the normalization procedure outlined in scDCC [16]. Specifically, a Standard Scaler was applied, ensuring that each cell has a mean of 0 and a standard deviation of 1.

For the graph-based algorithm, instead of performing a direct dimensionality reduction on the initial data, we selected the 5000 genes with the highest upward deviation from the adjusted variance. To achieve this, the variance *σ*^2^ and mean *μ* for each gene in the log-normalized count matrix are calculated. Using these values and following the same idea as presented in [32], we fitted a curve of the form

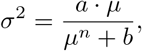

finding the best possible values of *a* and *b*. The selected genes are the 5000 genes with the largest difference between the actual variance and the expected variance.

### 3.4 Metrics

To evaluate the performance of the proposed models, we used three different supervised metrics. Specifically, we used accuracy, normalized mutual information (NMI), and the adjusted Rand index (ARI) to assess the degree of correspondence between the clusters identified by the algorithm and the true clusters.

#### 3.4.1 Accuracy

Accuracy for clustering is defined as the percentage of correctly assigned clusters within the dataset. If *n* is the number of data points, *y* represents the true clusters, and *ŷ* denotes the clusters assigned by the model, then accuracy is defined as

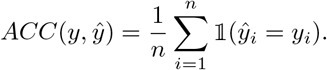

A contingency or confusion matrix is computed to establish correspondence between labels, and labels are rearranged to find the best match between cluster labels and true labels. Based on this correspondence, an accuracy score can then be calculated.

#### 3.4.2 Normalized Mutual Information (NMI)

NMI measures the mutual information between predicted clusters and ground-truth labels. It ranges from 0 to 1, where 1 indicates perfect clustering. If *K* is the number of clusters and *U* = *{U*_1_, *U*_2_, …, *U*_*K*_*}* and *V* = *{V*_1_, *V*_2_, …, *V*_*K*_*}* are the predicted clusters and ground-truth labels, the mutual information between *U* and *V*, denoted as *I*(*U, V*), is defined as

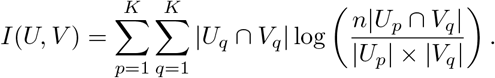

The entropies of a clustering assignment are defined as

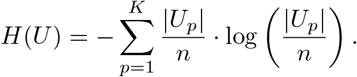

Finally, the **NMI** metric is defined as

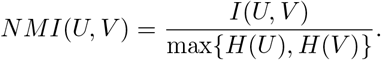

#### 3.4.3 Adjusted Rand Index (ARI)

ARI compares the similarity between clustering results and ground-truth labels. It ranges from *−*1 to 1, calculated as

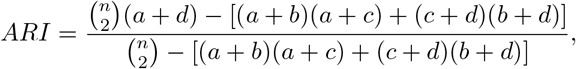

where *a* is the number of pairs in the same group in both *U* and *V*; *b* is the number of pairs in different groups in both *U* and *V*; *c* is the number of pairs in the same group in *U* but in different groups in *V*; and *d* is the number of pairs in different groups in *U* but in the same group in *V*.

### 3.5 Tools

We compared the supervised metrics produced by our algorithms to those obtained using three of the most widely used tools for scRNA-seq data clustering: Seurat [10], SC3 [6], and scDCC [16]. Each of these tools employs different clustering methodologies. Seurat uses a graph-based clustering algorithm on the *k*-NN graph of the cells in a lower-dimensional representation. SC3 utilizes a *k*-means algorithm combined with a consensus hierarchical clustering approach to determine the final clusters, requiring the number of clusters to be specified beforehand. Finally, scDCC is a deep neural network that follows an autoencoder architecture, modeling the dataset with a zero-inflated negative binomial distribution as described in [15]. It includes a novel clustering loss function based on the Kullback-Leibler (KL) divergence, which aids in computing the clusters directly within the neural network.

For Seurat and SC3, we followed the procedures outlined in their tutorials using the default parameters. In the case of scDCC, we had to make several modifications to the code in order to make it work with our current hardware architecture, and to fix bugs in the code. In particular, the script tried to read files that were either missing from the repository or unrelated to the scRNA-seq experiment data.

## 4 Validation

Figure 3 shows the distribution of accuracy values achieved by the two methods developed in this work, compared to Seurat, SC3, and scDCC. Detailed results for the ARI and NMI metrics are provided in the supplementary Figures 1 and 2. In the analysis of 24 simulated datasets, SC3 and scDCC demonstrated the greatest stability, consistently achieving higher supervised metric values across the different methodologies, with an overall mean accuracy of 90% in both methods. The deep learning approach AE-GMM achieved the third best mean accuracy with a value of 88.3%, followed by the *k*-MST algorithm with a mean accuracy of 86%. Finally, Seurat showed the lowest performance with a mean accuracy of 80%. It is important to note that these last two algorithms do not receive the number of clusters as an input parameter, which explains their position in the ranking of methods.

**Fig. 3:**
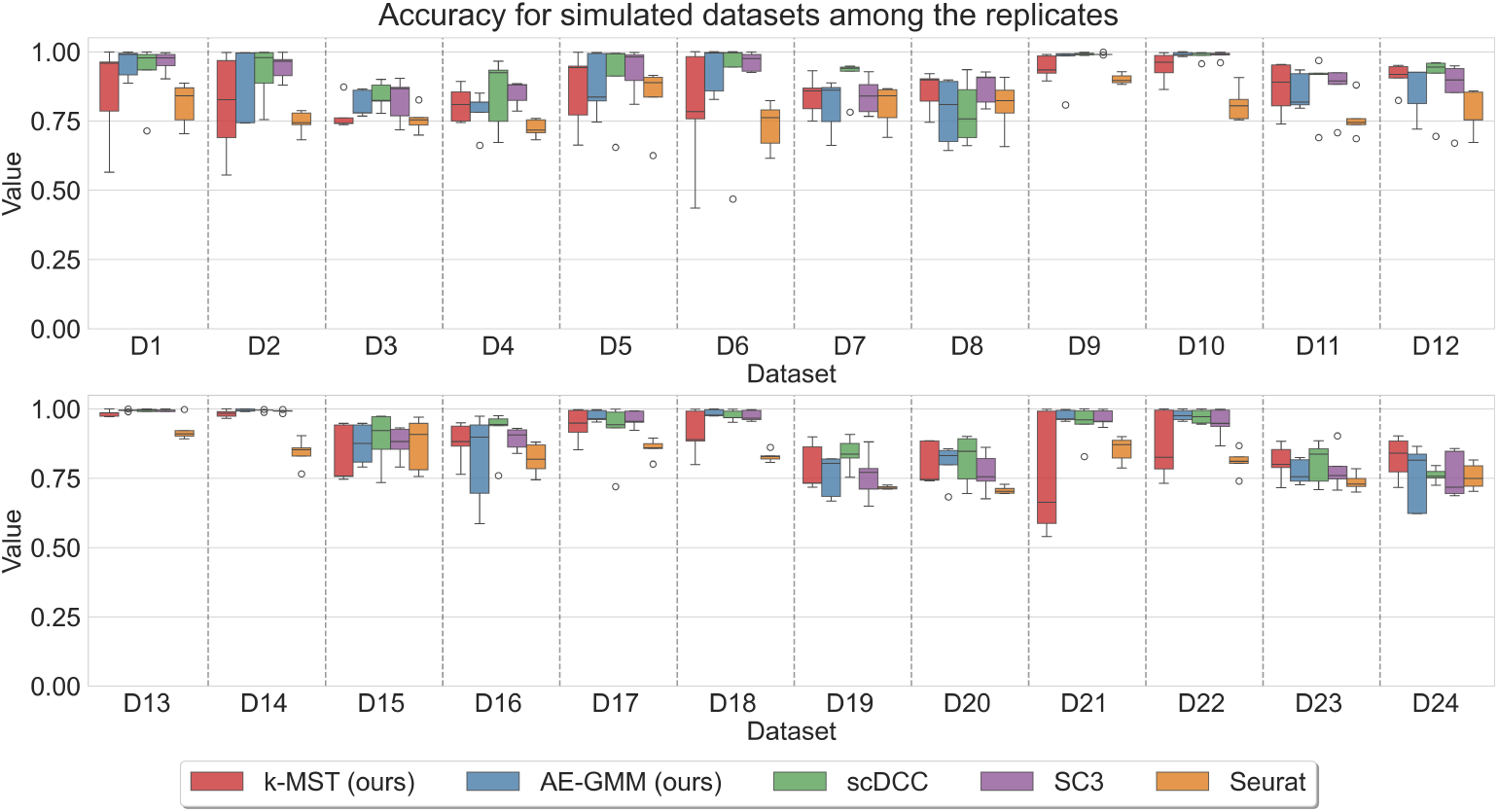
Comparison of accuracy across different methods on simulated datasets. The figure presents the accuracy of six methods (k-MST, AE-GMM, scDCC, SC3 and Seurat) evaluated on 24 simulated datasets (D1 to D24). The values are represented as boxplots, illustrating the variability in accuracy among replicates for each dataset.

AE-GMM achieved the highest metrics across replicates in seven datasets: D10, D13, D14, D17, D18, D21, and D22. All of these datasets contained only 3 clusters. In contrast, the *k*-MST algorithm achieved the best performance in datasets D12, D23, and D24. Each of these datasets contained 5 clusters.

Figure 4a presents the accuracy results for the publicly available datasets. The deep learning approaches were executed 10 times for each dataset with different seeds to mitigate the dependency of the results on the random initialization of weights. In contrast with the simulation experiments, the SC3 had the less stable performance with a reduction of over 10% compared to the other methods in the Human Liver and Mouse Retina datasets. In particular for the Mouse Retina dataset, the AE-GMM aproach presented in this work achieved an increase in accuracy over 5% compare to previous approaches. The AE-GMM ranked first in this dataset and in the Worm Neuron dataset. In the PBMC dataset, the scDCC method delivered the highest performance, while AE-GMM, along with SC3, tied for second-best results. However, AE-GMM ranked fourth in the Human Liver dataset. In this dataset, the *k*-MST method ranked first, outperforming the other tools by more than 8%, followed by the scDCC algorithm. The same behavior is presented in the Mouse Bladder dataset, where the *k*-MST algorithm obtained the best accuracy. In contrast with the results observed in the simulations, the behavior of this method was stable in these experiments, improving the accuracy of previous methods in two of the five datasets, and surpassing the Seurat results in three of the five experiments. Overall, the *k*-MST methodology achieved a mean accuracy of 68.4% across all datasets, and the AE-GMM achieved an average of 68.5%. Both averages were higher than those obtained by Seurat (65.4%), SC3 (57.9%), and scDCC (68.3%). The NMI and ARI results are shown in the supplementary figures 3 and 4.

Figure 4b shows a comparison of the accuracy of different preprocessing steps for the *k*-MST algorithm. The mean-variance approach uses the gene selection method described in Section 3.3. In the only-variance approach, the 5000 genes with the highest variance were selected prior to computing cell correlations. The no filter approach incorporates all the genes in the dataset without filtering. Additionally, we evaluated the impact of dropout correction using the method proposed in [15], where an autoencoder with ZINB loss was employed to address dropouts. The output of this neural network was treated as a corrected dataset and used as input for the algorithm, both with the selection of the top 5000 genes (mean-variance) and without gene selection (no filter).

**Fig. 4:**
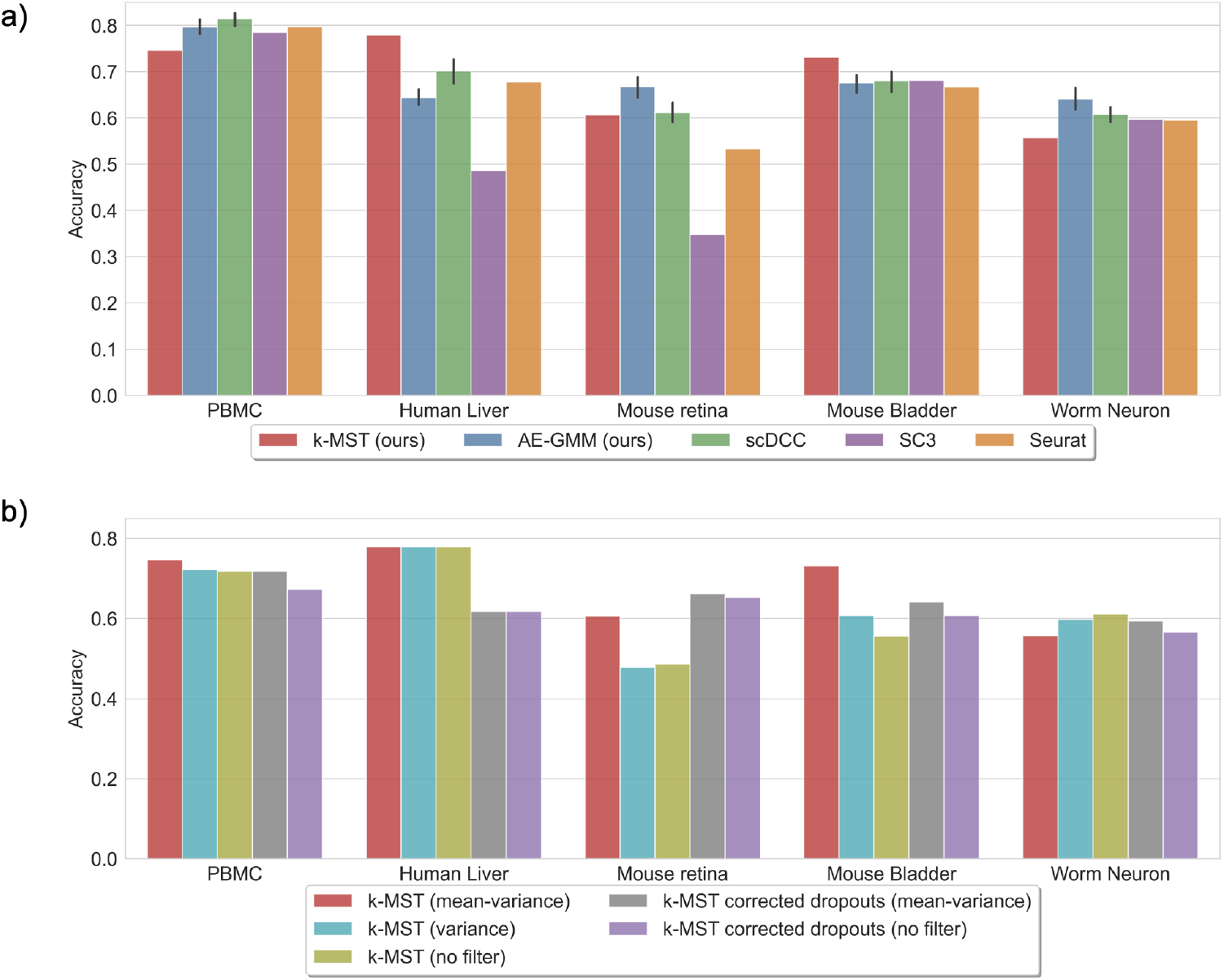
Comparison of accuracy across different methods on open-source datasets. a) Proposed methods against scDCC, SC3 and Seurat. b) Comparison of the *k*-MST approach for different preprocessing techniques.

In the PBMC dataset, all methods showed similar performance, with an average accuracy of 71%. The best-performing algorithm for this dataset was the meanvariance gene selection approach. For the Human Liver dataset, the methods without dropout correction produced better results, showing an improvement of over 16% compared to the other techniques. As this dataset contained only 5000 genes, the various gene selection algorithms yielded identical outcomes. In the Mouse Retina dataset, the dropout correction method outperformed the other approaches, followed by the mean-variance selection algorithm. This was the only dataset where such behavior was observed. In the Mouse Bladder dataset, the mean-variance method outperformed the others by 9%, followed by the same gene selection method with dropout correction. Finally, in the Worm Neuron dataset, all methods performed similarly, with a slight advantage for the no filter approach.

### 4.1 A new dataset for benchmarking of scRNA-seq cell clustering methods

Single nuclei RNA sequencing of the epilepsy patient resulted in a total of 1, 137 × 10^6^ reads. From these, 97.4% mapped to the human reference genome with 70.2% of reads mapping uniquely to an intronic region. The raw count matrix had a total of 91.29% zero values before gene filtering. After filtering cells with abnormal mitochondrial content, cells with an aberrant number of expressed genes, and doublets the dataset included a total of 4,018 high-quality cells having expected values for the number of molecules per cell, the number of genes per cell and the mitochondrial gene content for cells. The final distribution of counts per cell and sequenced genes are presented in Figure 5 a.

**Fig. 5:**
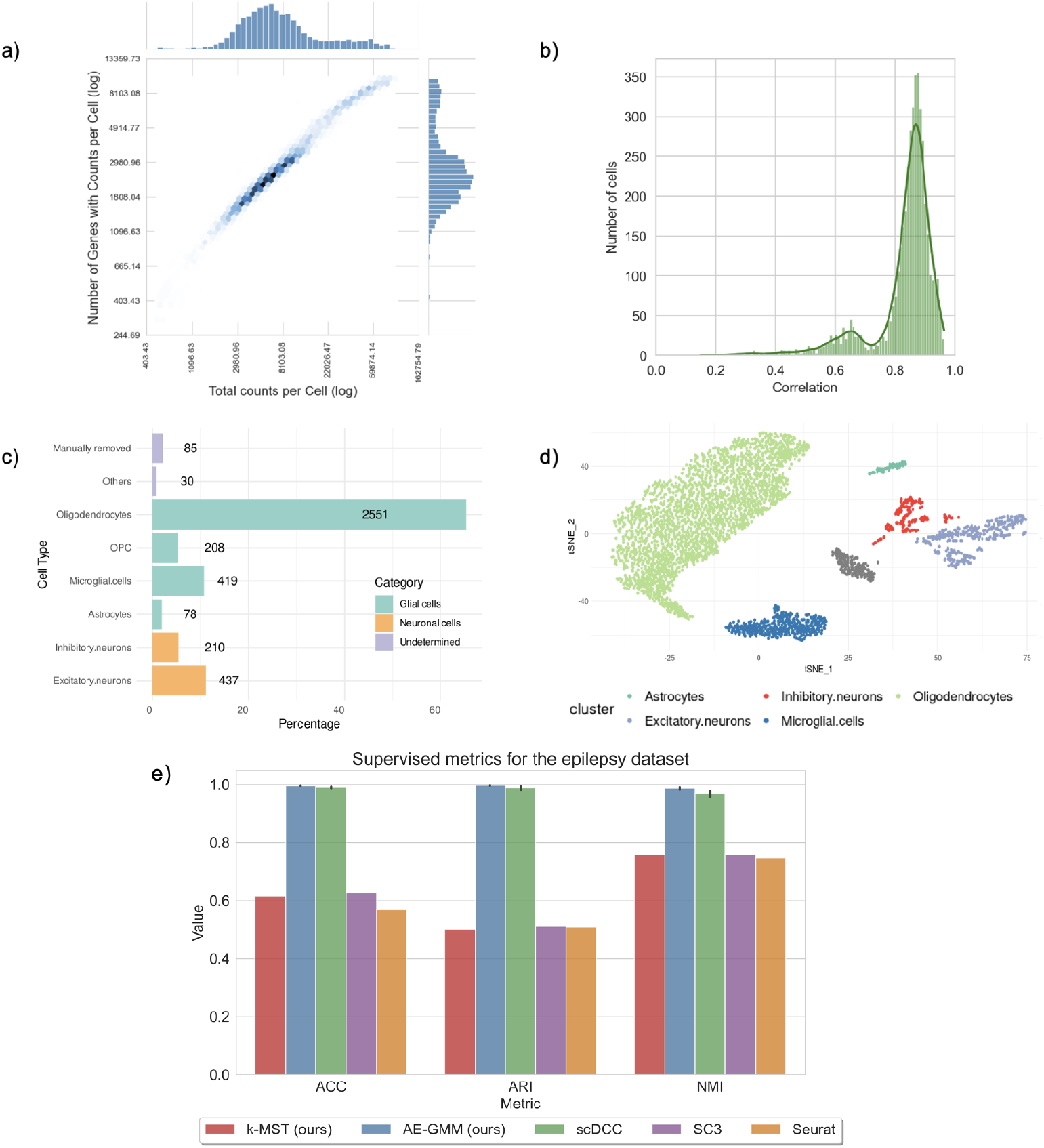
a) Sequencing depth of the sample. b) Histogram of maximum correlation values between cells and clusters defined at the human protein Atlas. c) Percentage values of cell types presented in Epilepsy dataset. The count of cells for each type is indicated. d) t-SNE visualization of cell types used as the epilepsy gold standard dataset. e) Supervised metrics for the epilepsy dataset among the different methodologies.

Correlations between the gene profiles of the sequenced cells and the Human Protein Atlas [33] database were calculated to perform the supervised assignment method described above. Figure 5b shows that the maximum correlations were distributed between 0.7 and 1 for most cells, with a mean value of 0.83. As shown in Figure 5c, the dataset was dominated by Oligodendrocytes, representing 63.9% (n=2552) of the cells, followed by Excitatory cells at 11.7% (n=467), and Microglial cells at 11.3% (n=452). With less than 300 representatives, the Inhibitory Neurons represent 5.56% (n=222), followed by oligodendrociyte Precursor Cells (OPC) with 5.34% (n = 213), and a final percentage of 2.05% (n=82) for Astrocytes. Figure 5d shows a t-SNE visualization of the annotated cells. Based on this analysis, 85 probably misclassified cells (42 neurons, and 43 glial cells) were removed from the dataset for benchmarking purposes. A final number of 3,903 annotated cells were used as the gold standard after removing the “Others”category.

After assigning cell types based on the Human Protein Atlas database, we executed the proposed algorithms and used the assigned clusters to compute the three supervised metrics. In this case we also ran SC3, Seurat and scDCC and compared the results. As shown in Figure 5, the AE-GMM methodology achieved the highest scores across all three metrics, with an average accuracy of 95.2% and a standard deviation of 0.7% over 10 executions of the algorithm. The second-best performance was achieved by scDCC with a small difference in metrics compared to AE-GMM. SC3, Seurat and *k*-MST showed similar results across the three supervised metrics. These results suggest that the highest-performing methodologies for the epilepsy dataset are those based on neural networks.

Supplementary Figure 5 presents the t-SNE representation of the clustering techniques applied to the epilepsy dataset. The clustering methods SC3, *k*-MST, and Seurat exhibit overclustering within the oligodendrocyte cells, dividing it into two or three subclusters. Adjusting the Seurat results by merging clusters 0 and 1 leads to enhanced clustering performance, with an accuracy of 88.4%, ARI of 97.3%, and NMI of 89.1%. For *k*-MST, merging clusters 0, 2, and 10 results in an accuracy of 94.1%, NMI of 93.2%, and ARI of 97.1%. Similarly, combining clusters 1 and 2 for SC3 yields an accuracy of 92.1%, NMI of 92.3%, and ARI of 96.9%. These findings indicate that these methods benefit from post-processing adjustments to correct for overclustering. The improvement in clustering metrics after merging suggests that such adjustments yield a more biologically accurate representation of cell populations in the dataset, showing the best results in the *k*-MST approach.

## 5 Discussion

Identification of cell types from scRNA-seq data is an interesting bioinformatics problem, due to the high dimensionality and non linearity of the data. Given the wide use of scRNA in different experimental settings in a large number of species, software implementing accurate algorithms for unsupervised cell clustering of scRNA data is likely to have a large number of users. In this work we contributed two algorithmical improvements building upon two major strategies for scRNA cell clustering (graph based and autoencoders). Experiments with a large number of datasets show that none of the methods is inherently superior, but that the proposed methods achieve good results in more scenarios compared to previous approaches. Both methods are implemented in the software package scRANGE, to facilitate independent reproduction of the results and to facilitate the use of these methods by different research groups. Our experience with current methods for cell clustering in scRNA data suggests that implementation using basic software engineering practices is important to translate good algorithmic ideas to useful solutions. Although a large number of algorithmic approaches for clustering single-cell data have been published in recent years, Seurat is a current standard de-facto for the analysis of these data, primarily due to the usability of the tool. Conversely, the potential improvement in accuracy that could be achieved using models such as that implemented in scDCC can not be realized due to software bugs and issues with version management.

The *k*-MST algorithm proposed in this article is a graph-based clustering method. It primarily differs from Seurat in two key aspects: Firstly, it does not employ a dimensionality reduction technique such as PCA; instead, it performs gene selection. Secondly, the graph used (*k*-MST) differs from the (*k*-NN) graph implemented in Seurat. Although in both graphs each node has at least *k* neighbors, the *k*-MST allows more edges per node for nodes with a large number of low cost connections. We believe that this allows to incorporate more relational information between cells, improving the outcome of the subsequent clustering using the Louvain algorithm. An advantage of this method over methods based on autoencoders is that it does not require prior knowledge of the number of clusters to be generated. The Louvain algorithm identifies the optimal number of clusters without the need for preliminary manual analysis.

The second proposed algorithm, AE-GMM, is a Deep Learning-based method. Unlike traditional autoencoder approaches, this algorithm integrates clustering directly into the training process, learning the parameters that characterize each cluster. The inclusion of the GMM loss allows the resulting clusters to have more flexible shapes, such as ellipsoids, in contrast to methods like *k*-Means, which impose stricter constraints on cluster shapes. This flexibility allowed the algorithm to obtain the highest supervised metrics on three of the six datasets considered in this study. However, this method shares a common limitation with many other clustering techniques (such as SC3, scDCC and k-Means), which is the need to know the number of clusters in advance to define the network parameters before training.

Finally, this article presents a new single-cell RNA-seq dataset obtained from a patient with refractory epilepsy. Thanks to the decreasing costs of sequencing, this dataset was sequenced with a depth per cell higher than that obtained in previous datasets used for benchmarking of scRNA-seq clustering methods. This translated into a lower percentage of zero entries. We believe that this approach will be taken by different groups currently designing scRNA-seq experiments. Moreover, we used a supervised approach to build the gold standard cell annotations. Although we acknowledge that this method could produce some erroneous assignments, the proposed method is orthogonal to the unsupervised clustering methods. Hence, errors should not be biased to favor any particular clustering algorithm. We believe that this dataset can contribute to increase the heterogeneity of benchmark datasets, providing a useful resource for developers of scRNA-seq data analysis methods.

## 6 Data and software availability

The algorithms and the data sets supporting the results presented in this article are publicly available in github as part of the distribution of *scRANGE*.

## 7 Declarations

### 7.1 Ethical Approval

The epilepsy sample was obtained from a pediatric patient at HOMI, through a collaborative project among the institutions involved in this manuscript. The project was approved by the research ethics committee at HOMI, as evidenced by the minute 49-21 of the meeting held on June 1st of 2021. The brain tissue was obtained as part of a prescribed surgery procedure for the patient, following all applicable guidelines and regulations. Legal representatives of the patient signed an informed consent for research purposes and data were anonymized to avoid identification of the patient.

### 7.2 Consent for publication

Not applicable

### 7.5 Author’s Contributions

JD conceived the study. LG, PS, JGP, DGO and AN collected the samples. JDR, PS and SM performed lab work. MR, CF, and JD developed software. MR, JDR, SO and AQ performed data analysis. LG, AN, SM, AQ and JD coordinated the work and provided scientific support. MR, JDR and JD wrote the manuscript. All authors read and approved the final version of the manuscript.

## 8 Acknowledgements

We acknowledge the high-performance computing unit of Universidad de Los Andes for their technical support to conduct the benchmark experiments presented in this manuscript.

