## Supplementary figures for "New algorithms for unsupervised cell clustering from scRNA-seq data"

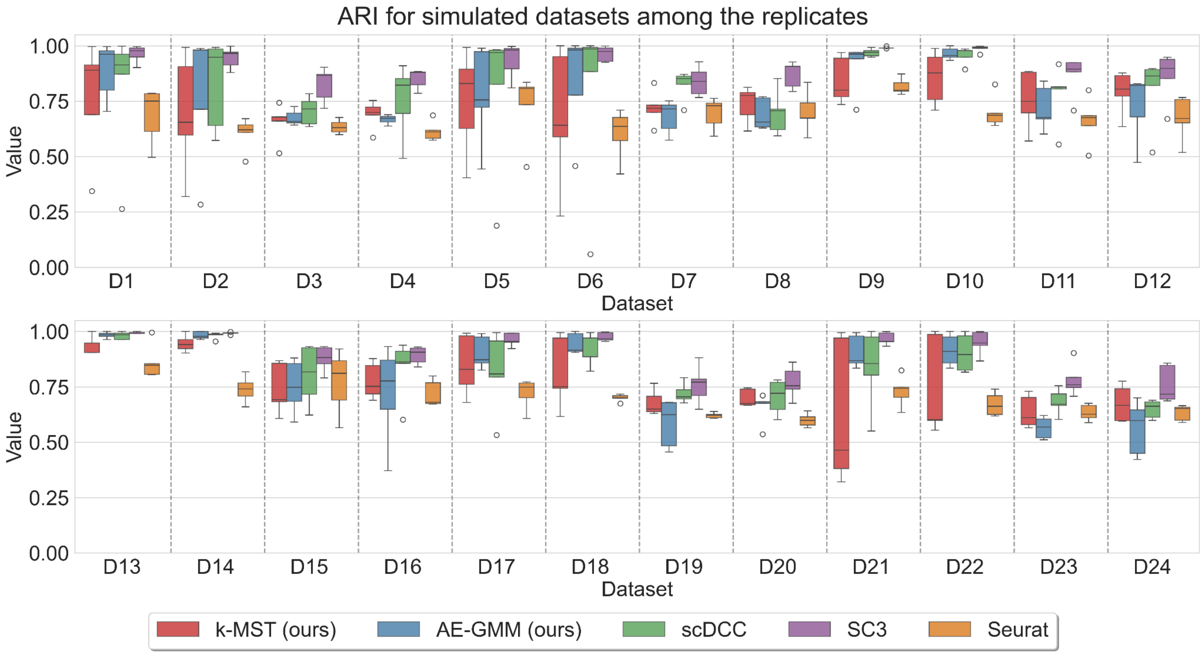


**Figure 1**. Comparison of ARI across different methods on simulated datasets. The figure presents the ARI of six methods (k-MST, AE-GMM, scDCC, SC3 and Seurat) evaluated on 24 simulated datasets (D1 to D24). The values are represented as boxplots, illustrating the variability in ARI among replicates for each dataset.

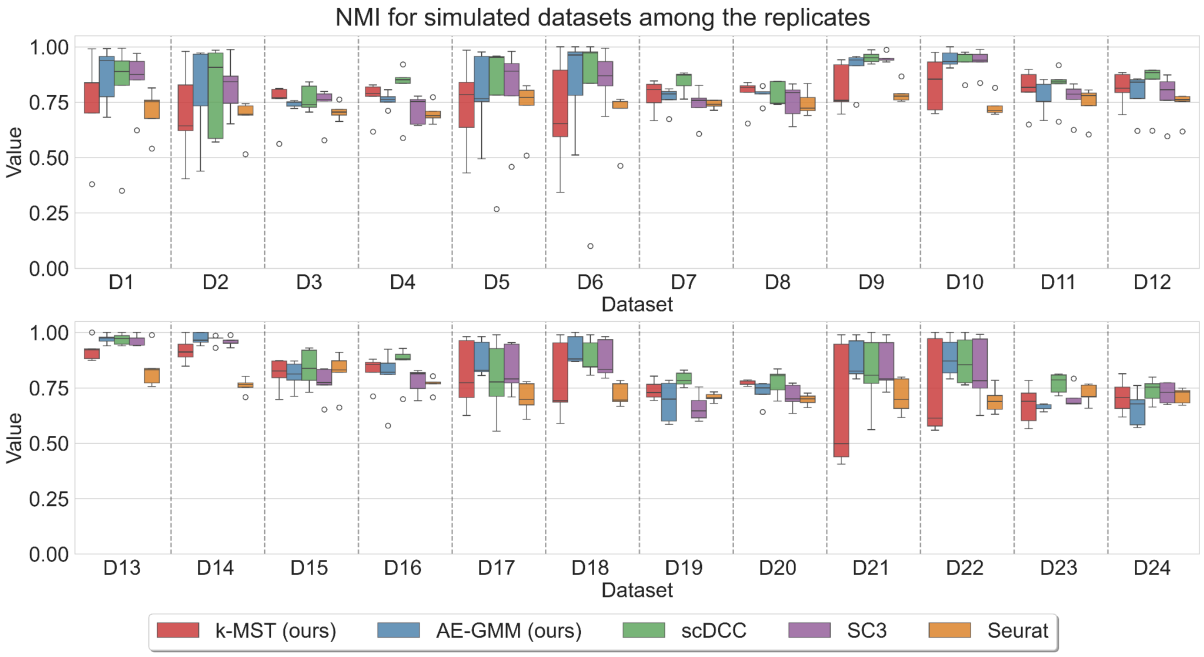


**Figure 2**. Comparison of NMI across different methods on simulated datasets. The figure presents the NMI of six methods (k-MST, AE-GMM, scDCC, SC3 and Seurat) evaluated on 24 simulated datasets (D1 to D24). The values are represented as boxplots, illustrating the variability in NMI among replicates for each dataset.

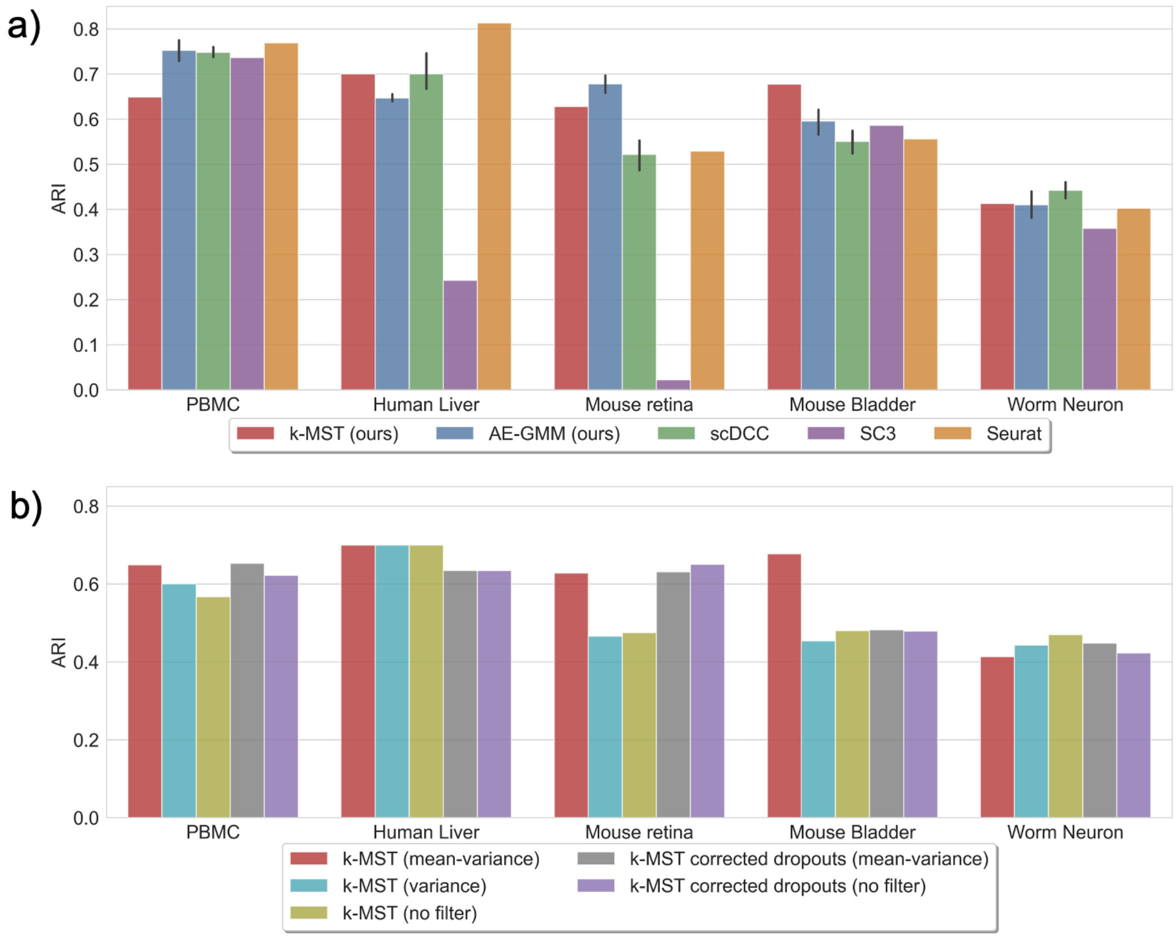


**Figure 3:** Comparison of ARI across different methods on open-source datasets. a) Proposed methods against scDCC, SC3 and Seurat. b) Comparison of the k-MST approach for different preprocessing techniques.

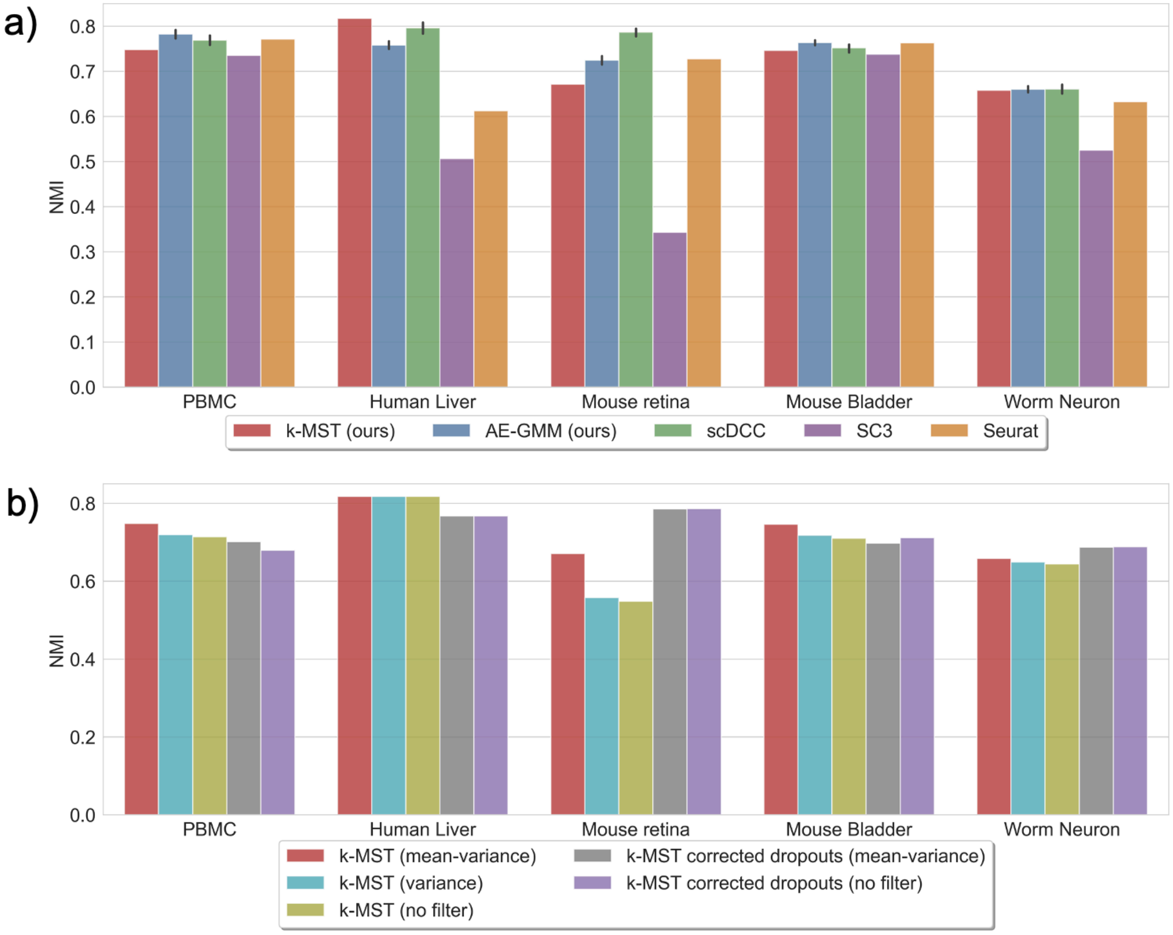


**Figure 4:** Comparison of NMI across different methods on open-source datasets. a) Proposed methods against scDCC, SC3 and Seurat. b) Comparison of the k-MST approach for different preprocessing techniques.

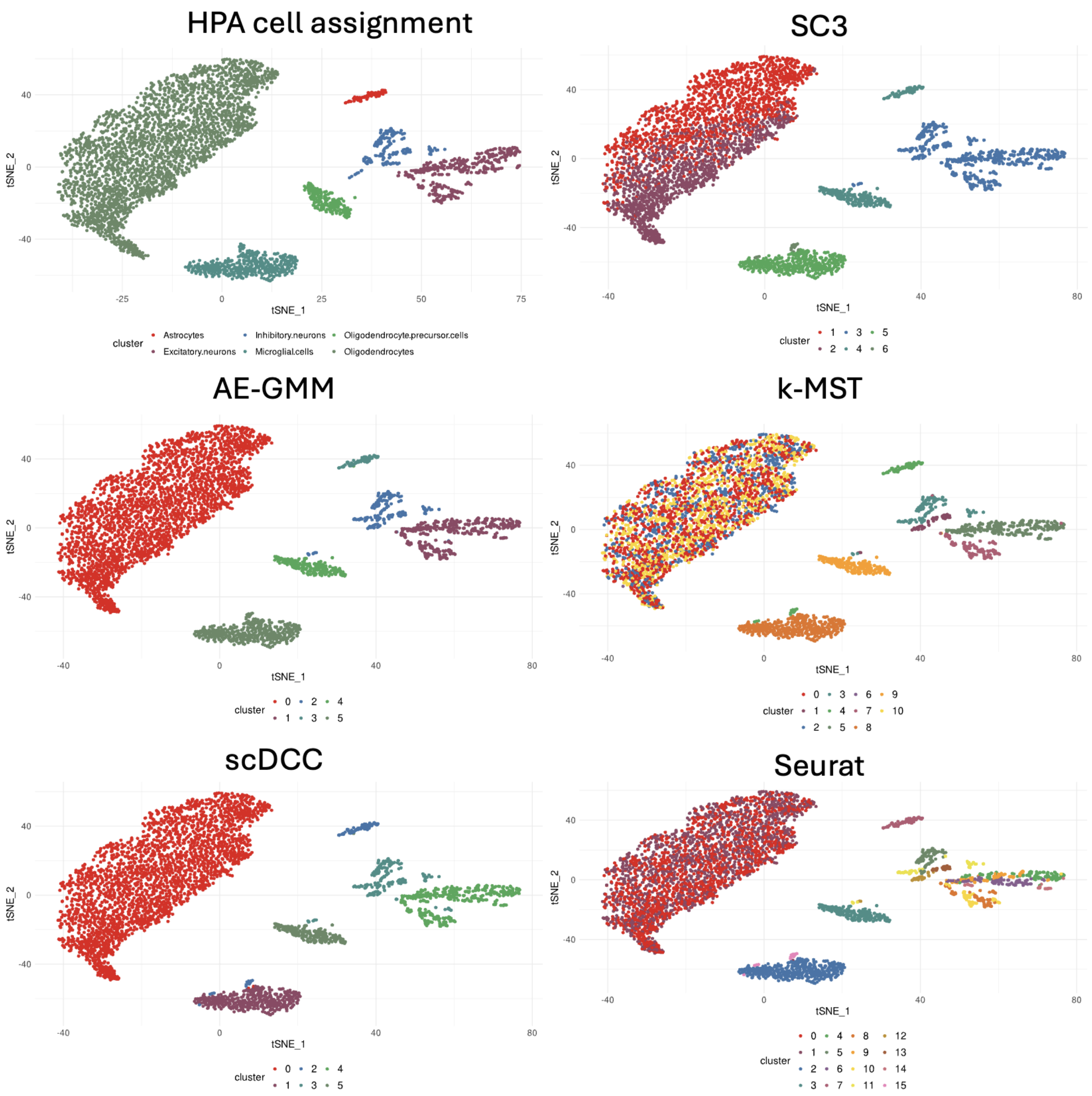


**Figure 5:** t-SNE representation of the results of different clustering techniques for the epilepsy dataset.
